# Single-Cell Multimodal Profiling Reveals a Novel CD26^+^ Fibroblast Subpopulation in Atherosclerosis

**DOI:** 10.1101/2025.01.27.635067

**Authors:** Alexander C. Bashore, Johana Coronel, Chenyi Xue, Lucie Y. Zhu, Muredach P. Reilly

## Abstract

**Background:** Atherosclerosis involves complex interactions between lipids, immune cells, vascular smooth muscle cells (VSMCs), and fibroblasts within the arterial wall. While significant advances in single-cell technologies have shed light on the roles of immune cells and VSMCs in plaque development, fibroblasts remain underexplored, leaving critical gaps in understanding their contributions to disease progression and plaque stability. Comprehensive characterization of fibroblast phenotypes in atherosclerosis is essential to unravel their diverse functions and to distinguish between subsets that may play protective versus pathogenic roles in the disease process.

**Methods:** Here, we utilized CITE-seq (Cellular Indexing of Transcriptomes and Epitopes by Sequencing) to comprehensively profile fibroblast diversity in a mouse model of atherosclerosis. Mice were fed an atherogenic diet for 0, 8, 19, and 26 weeks, representing distinct stages of disease progression, enabling a detailed phenotypic characterization of fibroblasts throughout the course of atherosclerosis development.

**Results:** We identified four distinct fibroblast subpopulations, including a myofibroblast population closely resembling VSMC-derived chondromyocytes. The proportions of these fibroblast subsets exhibited a modest decline as atherosclerosis progressed. Through multimodal analysis, we identified CD26 as a highly expressed and specific marker for one of these fibroblast subpopulations, distinguishing it from other subsets. Using a combination of flow cytometry and immunohistochemistry, we demonstrated that CD26^+^ fibroblasts predominantly reside in the adventitia of healthy arteries. During atherosclerosis progression, these cells expand into the intima and primarily localize within the fibrous cap of the lesion.

**Conclusions:** Our multi-omic analysis highlights the phenotypic diversity and dynamic changes of fibroblasts during atherosclerosis progression. Among these, CD26+ fibroblasts emerge as a distinct subpopulation that expands within atherosclerotic lesions and may play a critical role in promoting plaque stability through their migration into the fibrous cap.

## Introduction

Atherosclerosis, a leading cause of cardiovascular diseases, involves complex pathophysiological processes characterized by the accumulation of lipids, inflammatory cells, and fibrotic cells within the arterial wall^1^. Atherosclerotic plaques are composed of a complex mixture of immune cells, endothelial cells, vascular smooth muscle cells (VSMCs), and fibroblasts. The walls of arteries exhibit a heterogeneous structure comprised of three distinct layers: the tunica intima, tunica media, and tunica adventitia^2^. Each layer possesses unique histological, biochemical, and functional characteristics, playing distinct roles in maintaining vascular homeostasis and regulating vascular responses to stress and injury. Identifying the origin and phenotypic diversity of cells within atherosclerotic plaques is crucial to understanding the causal and mechanistic roles of newly described cell types.

Recent advancements in single-cell technologies have significantly enhanced our ability to characterize the cellular landscape of human^3–6^ and mouse^7–13^ atherosclerosis. These tools have been instrumental in identifying cell type-specific genes and molecular mechanisms, providing insights for novel therapies. Most studies have focused on vascular smooth muscle cells (VSMCs) and immune cells, revealing their roles in plaque development and inflammation. However, research on fibroblasts has been limited, with only one study applying these technologies to explore their biology in atherosclerosis^14^. Given the well-documented plasticity and heterogeneity of fibroblasts in other tissues^2,15^, further investigation is needed to identify and characterize distinct fibroblast subsets in atherosclerosis and elucidate their biological relevance. These efforts could uncover how fibroblasts contribute to plaque formation, progression, and stability, potentially revealing new therapeutic targets.

In this study, we utilized CITE-seq (Cellular Indexing of Transcriptomes and Epitopes by Sequencing) to perform a comprehensive phenotypic characterization of fibroblast subpopulations within atherosclerotic plaques in mice. This cutting-edge approach allowed us to not only capture gene expression profiles but also integrate protein marker data, providing a multi-omic perspective on fibroblast heterogeneity in atherosclerosis. Our analysis uncovered an unprecedented diversity of fibroblast subpopulations, shedding light on the complexity of fibroblast involvement in plaque development. Among the diverse subsets identified, we focused on a distinct CD26^+^ fibroblast subpopulation. Importantly, we were able to isolate and spatially map this subset within atherosclerotic lesions. Our findings revealed that this CD26^+^ fibroblast population undergoes dynamic change over time on a western diet (WD) and lesion progression being present in the adventitial layer at baseline before WD and lesion formation and found extensively in the lesion intima as disease advances on WD. This suggests an active role for these fibroblasts in the structural remodeling of vessel and intima during progression of the plaque and highlights their potential involvement in the regulating plaque functions including lesion stability. Our study provides new insights into fibroblast biology in atherosclerosis and opens avenues for investigating how distinct subsets of fibroblasts contribute to the pathogenesis and progression of the disease.

## Methods

### Mouse studies

All animal experiments were conducted in compliance with the Institutional Animal Care and Use Committee (IACUC) of Columbia University under protocol number AABQ5576. Mouse strains were obtained from The Jackson Laboratory, including Ldlr KO (Stock: 002207), ZsGreen (Stock: 007906), and Myh11-CreERT2 (Stock: 019079). These strains were bred on a C57BL/6J background to create a hyperlipidemic smooth muscle cell lineage tracing model: Ldlr KO; ZsGreen; Myh11-CreER^T28^. Only male mice were used due to the insertion of the Myh11-CreERT2 transgene on the Y chromosome. To activate Cre recombinase and enable lineage tracing with ZsGreen, mice received five intraperitoneal tamoxifen injections (40 mg/kg body weight). Following tamoxifen administration, a minimum washout period of ten days was observed before commencing any experimental analyses. To induce atherosclerosis, mice were placed on a high-fat, high-cholesterol diet for defined durations (Research Diet: D12079Bi).

### Aorta digestion for CITE-seq preparation

At sacrifice, the vasculature was immediately perfused with ice-cold DPBS containing 1 µg/mL Actinomycin D (Thermo: 11805017). The aorta, including the ascending aorta, brachiocephalic artery (BCA), and thoracic aorta, was harvested and digested in RPMI with a mixture of 4 U/mL Liberase (Roche: 05401119001), 60 U/mL Hyaluronidase (Millipore Sigma: H3506-100MG), and 60 U/mL DNase (Worthington: LS006331). The resulting cell suspension was washed once with FACS buffer (2% HI-FBS, 5 mM EDTA, 20 mM HEPES, 1 mM Sodium Pyruvate in 1X DPBS) and centrifuged at 400g for 5 minutes at 4°C. Cells were then resuspended in 49 µL of FACS buffer, and 1 µL of TruStain FcX PLUS (anti-mouse CD16/32) (BioLegend: 156603) was added for 10 minutes at 4°C to block Fc receptors. The TotalSeq-A Mouse Universal Cocktail (BioLegend: 199901) oligo-conjugated antibodies were reconstituted in 100 µL of FACS buffer, and 30 µL of this cocktail was added to each single-cell suspension, incubating for 30 minutes at 4°C. Samples were washed twice in FACS buffer with centrifugation at 400g for 5 minutes at 4°C. For cell hashing, pellets were resuspended in Cell Multiplexing Oligos (10x Genomics) and incubated for 5 minutes at room temperature, followed by three washes in FACS buffer with centrifugation at 400g for 5 minutes at 4°C. Single-cell suspensions were stained with DAPI (1:3,000) and DRAQ5 (1:1,000). Viable cells, defined as DRAQ5-positive and DAPI-negative, were sorted using a BD FACSAria II and collected into 1.5 mL Eppendorf tubes containing DMEM/F12 + 10% FBS. Cells were then pelleted and counted for input into the 10x Genomics Chromium system.

### CITE-seq library preparation

CITE-seq libraries were prepared as previously described^6,9,16^. We adhered to the 10x Genomics 3’ v3 protocol per the manufacturer’s instructions for cDNA amplification using 0.2 mM of ADT additive primer (5’CCTTGGCACCCGAGAATTCC). The supernatant from the 0.6x SPRI cleanup was saved and purified with two rounds of 2x SPRI, and the final product was used as a template to produce ADT libraries. Antibody tag libraries were generated by PCR using Kapa Hifi Master Mix (Kapa Biosciences KK2601), 10 mM 10x Genomics SI-PCR primer (5’AATGATACGGCGACCACCGAGATCTACACTCTTTCCCTACACGACGCTC), and Small RNA RPIx primer (5’CAAGCAGAAGACGGCATACGAGATxxxxxxGTGACTGGAGTTCCTTGGCACCCGA GAATTCCA with X denoting one of the four following sequences: CGTGAT, ACATCG, GCCTAA, TGGTCA). Following amplification, Antibody tag libraries were cleaned with 1.6x SPRI. Subsequently, ADT quality was verified using a DNA high-sensitivity assay on an Agilent 2100 bioanalyzer.

### CITE-seq data pre-processing for mouse CITE-seq data

FASTQ files were processed using the Cell Ranger 6.1.2 pipeline, aligning sequencing reads to a custom mouse genome based on GRCm38 with the addition of the ZsGreen1 sequence^17^. Transcriptome mapping was performed against GENCODE M23 with ZsGreen1 annotation. For CITE-seq quantification, a reference containing 128 proteins, including 9 control antibodies, was incorporated. Separate count files were generated for each replicate (replicate 1, replicate 2, and replicate 3), and duplicate gene symbols from the GENCODE annotation were appended with numeric suffixes for distinction.

### Filtering and CarDEC_CITE clustering of mouse atherosclerosis progression CITE-seq data

Filtering and clustering of the CITE-seq data followed the approach described in Bashore et al.^6^. Briefly, RNAs and proteins were retained if they were present in at least 10 cells and 1 cell, respectively. Cell level filtering was performed using thresholds on feature and UMI count from RNAs and proteins. A maximum of 10% mitochondrial content was allowed in each cell. CarDEC_CITE was utilized for integration and clustering using 2500 highly variable RNAs and 30 highly variable proteins.

Differntial expression analysis was performed on each of the 22 clusters identified. Standard pipeline in Scanpy package was applied to RNA counts. Centered log ratio normalization from MUON package was applied to protein counts, followed by Wilcoxon rank sum test for differential expression. RNAs and proteins with FDR adjusted P-value <0.05 and presence in ≥25% cells in either group were considered differentially expressed.

### Flow Cytometry Analysis

At sacrifice, the vasculature was immediately perfused with ice-cold DPBS. The aorta, including the ascending aorta, BCA, and thoracic aorta, was harvested. To separate adventitia from media and intima, the whole aorta was incubated in Hank’s balanced salt solution (HBSS Ca^++^, Mg^++^) with a mixture of 175U/ml Collagenase II(Thermo Fisher Scientific, 17101015) and 1.25U/ml Elastase (Millipore Sigma, E7885) at 35°C for 15 min. After incubation, using fine forceps, the adventitia was peeled from the media and intima. Tissue was then digested that as with CITE-seq: RPMI with a mixture of 4 U/mL Liberase 60 U/mL Hyaluronidase, and 60 U/mL DNase. The resulting cell suspension was washed once with FACS buffer (2% HI-FBS, 5 mM EDTA, 20 mM HEPES, 1 mM Sodium Pyruvate in 1X DPBS) and centrifuged at 400g for 5 minutes at 4°C.Following digestions, cells were incubated with TruStain FcX PLUS (anti-mouse CD16/32) (Biolegend, 156603) for 10 minutes at 4°C to inhibit unspecific binding of antibodies to Fc receptors. Then cells were stained with fluorescently conjugated antibodies at 1:50 concentration for 30 min at 4°C: CD45-PerCPCy5.5 (BioLegend, 368503), CD31-BV711 (BioLegend, 102449), CD90.2-APC (BioLegend, 105312), and CD26-PE (BioLegend, 137804). Cell viability was assessed using DAPI. All samples were measured on a NovoCyte Penteon Flow Cytometer System, and the results were analyzed using Flowjo v10.10.0 software.

### Immunofluorescence

Hearts containing the aortic root were fixed in 4% paraformaldehyde for 24 hours, transferred to 70% ethanol, and embedded in paraffin wax. Paraffin blocks were sectioned at 6 µm using a Leica Microtome (RM2235) and floated on a 40°C water bath to ensure flat sections. Sections were mounted onto microscope slides (Fisher Scientific, 1255015), dried overnight, and stored at room temperature until staining or further analysis.Immunofluorescence was performed on paraffin sections of the aortic root. The sections were deparaffinized using xylene and 100% ethanol, followed by a Tris-based solution’s antigen retrieval process (Vector Laboratories, H3301250). Subsequently, sections were blocked with 10% normal goat serum for 1 hour to prevent nonspecific binding. Then, sections were incubated overnight at 4°C in a humidifier chamber with primary antibodies: ZsGreen (Origene, TA180002) and CD26 (Abcam, ab187048) at 1:200 concentration, along with IgG controls, which were obtained from Thermo-Fisher Scientific. The next day, the sections were washed with PBS and then incubated for 1 hour at room temperature with appropriate secondary antibodies. After additional washing with PBS, the sections were mounted with a ProLong Glass Antifade Mountant with NucBlue Stain (Thermo-Fisher Scientific, P36983). Imaging was conducted on a Nikon Eclipse Ti-S microscope, while image processing was performed using Image J.

## Results

### High-dimensional multi-omic profiling identified three distinct fibroblast subpopulations in atherosclerosis

To phenotypically describe fibroblasts within atherosclerotic lesions, we utilized our recently published CITE-seq dataset^9^, which characterized the expression of 119 cell surface proteins in mouse atherosclerosis. This dataset was derived from male LDLr^-/-^ ROSA26^LSL-ZsGreen1/+^Myh11-CreER^T2^ mice, where tamoxifen administration permanently induces ZsGreen1 expression in VSMCs. The mice were fed a WD for 0, 8, 16, and 26 weeks to represent the increasing severity of atherosclerosis progression. Our clustering analysis identified 22 distinct cell populations, including 4 fibroblast subpopulations (**Fig 1A**). Given the well-established phenomenon that VSMCs can acquire fibroblast-like characteristics during phenotypic modulation, we leveraged our VSMC lineage-traced mouse model to distinguish between VSMC-derived and non-VSMC-derived cells (**Fig 1B**). We identified three distinct fibroblast subpopulations (Fibroblast 1, Fibroblast 2, and Fibroblast 3) along with a myofibroblast population, which is partially composed of chondromyocytes derived from VSMCs^7,18^. To investigate the response of each fibroblast population to atherosclerotic conditions, we examined the proportion of each cluster throughout WD feeding (**Fig 1C**). In contrast to well-established increases in VSMC-derived cell types in lesions during progression of atherosclerosis^9^, this analysis revealed subtle decreases in the proportions of Fibroblast 2, Fibroblast 3, and myofibroblast, with no changes in Fibroblast 1. Additionally, we assessed the proportion of each non-VSMC-derived fibroblast subpopulation during the progression of atherosclerosis (**Fig 1D**). Fibroblast 1-3 populations each were consistently greater than 99% non-VSMC-derived. In contrast, the cluster that was composed of myofibroblasts and chondromyocytes was about 60% non-VSMC-derived at baseline, decreasing to 20% following 26 weeks of WD feeding.

**Figure 1.**
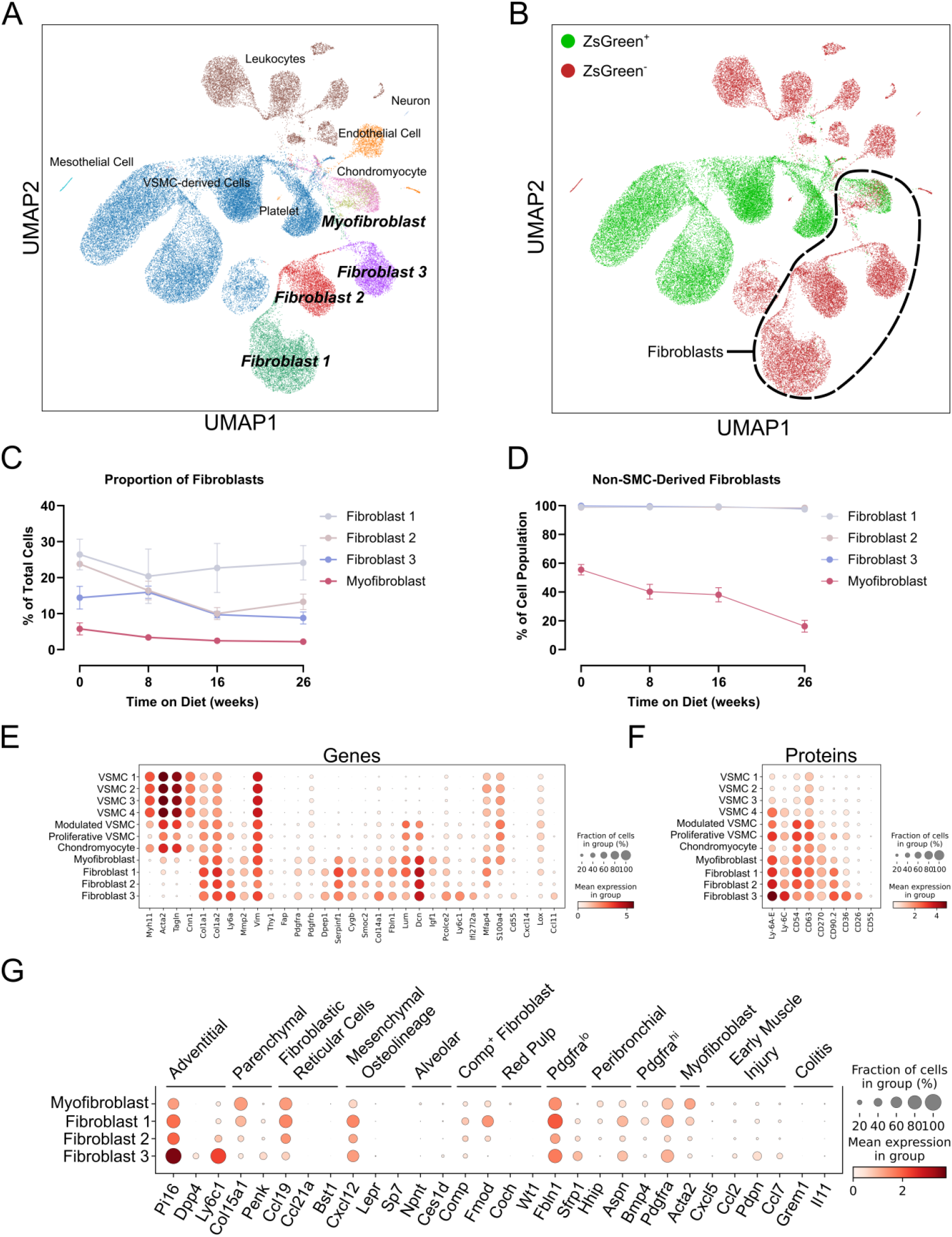
Multimodal analysis of fibroblast heterogeneity in atherosclerosis. (A) UMAP visualization of multimodal integration of all CITE-seq data (n=3 per time point) identified 22 distinct cell populations, highlighting 4 distinct fibroblast subpopulations. (B) UMAP categorizing cells by ZsGreen status. Circled cell clusters depict all fibroblast subpopulations. (C) Proportion of each fibroblast subpopulation throughout Western diet feeding. (D) Proportion of each fibroblast subpopulation that is ZsGreenthroughout Western diet feeding. (E) Dotplot depicting the expression of selected vascular smooth muscle cell (VSMC) and fibroblast genes across all VSMC and fibroblast clusters identified in our analysis. (F) Dotplot depicting top differentially expressed (DE) proteins across VSMC and fibroblast clusters. (G) Dotplot depicting expression of key genes from recent fibroblast atlas across our atherosclerotic fibroblasts.

To fully utilize our CITE-seq dataset, we assessed the expression of well-described fibroblast genes and newly identified cell surface proteins to identify distinguishing markers that could differentiate them from VSMC-derived cells. Our gene analysis confirmed that most fibroblast and VSMC genes lacked high specificity. Notably, some recently proposed specific genes, such as Pdgfra and Dpep1, were highly expressed in fibroblasts but were not uniformly expressed or were also expressed at lower levels in VSMC-derived populations. In our analysis, the most specific fibroblast-defining genes were Serpinf1 and Cygb (**Fig 1E**). Additionally, we identified CD36 and CD26 as subset-defining cell surface proteins for Fibroblast 3 (**Fig 1F**), distinguishing them from other fibroblast subsets. Interestingly, Fibroblast 3 displayed increased expression of Ly-6A/E compared to other fibroblasts, suggesting a role in proliferation and migration^19^.

To contextualize our identification of fibroblast subsets in atherosclerosis within other tissues and disease states, we analyzed the expression of key genes from a recently published comprehensive fibroblast atlas (**Fig. 1G**)^15^. This atlas described 10 fibroblast subsets at steady state and three additional subsets in perturbed states. Our findings revealed some shared characteristics between the fibroblast subsets identified in our study and those described in the atlas. Notably, Fibroblast 3 most closely resembled the adventitial pan-tissue fibroblast subset, marked by specific expression of genes such as Pi16, Dpp4, and Ly6c1. Myofibroblasts exhibited similarities to the parenchymal and myofibroblast subsets, suggesting a potential role in tissue remodeling and structural support. Fibroblast 1 displayed features of multiple subsets, including fibroblastic reticular cells, mesenchymal cells, red pulp fibroblasts, and COMP^+^ fibroblasts, reflecting a diverse functional capacity. In contrast, Fibroblast 2 did not align clearly with any defined subset in the atlas but appeared most closely linked to Pdgfra^lo^ fibroblasts, suggesting a potentially unique or undercharacterized fibroblast population. These comparisons highlight both conserved and tissue-specific features of fibroblast subsets in atherosclerosis.

### CD26^+^ Fibroblasts are functionally distinct and expand in the maturing lesion

To infer function difference between the four fibroblast subpopulations we identified, we used the top differentially expressed genes that defined each subpopulation (**Fig 2A**), we found a high degree of overlap, with each subpopulation showing upregulation of several collagen-related pathways. Since Fibroblast 3 appeared to be the most distinctive, we focused on uniquely upregulated pathways specific to this subset (**Fig 2B**). Interestingly, Fibroblast 3 exhibited upregulation of multiple pathways involved in lipid metabolism, including increased expression of Abca1 and Lpl. Sirtuin and Ferroptosis signaling pathways were also upregulated reflected in increased expression of Nfe2I2, Sod1, and Steap1 (**Fig 2C**).

**Figure 2.**
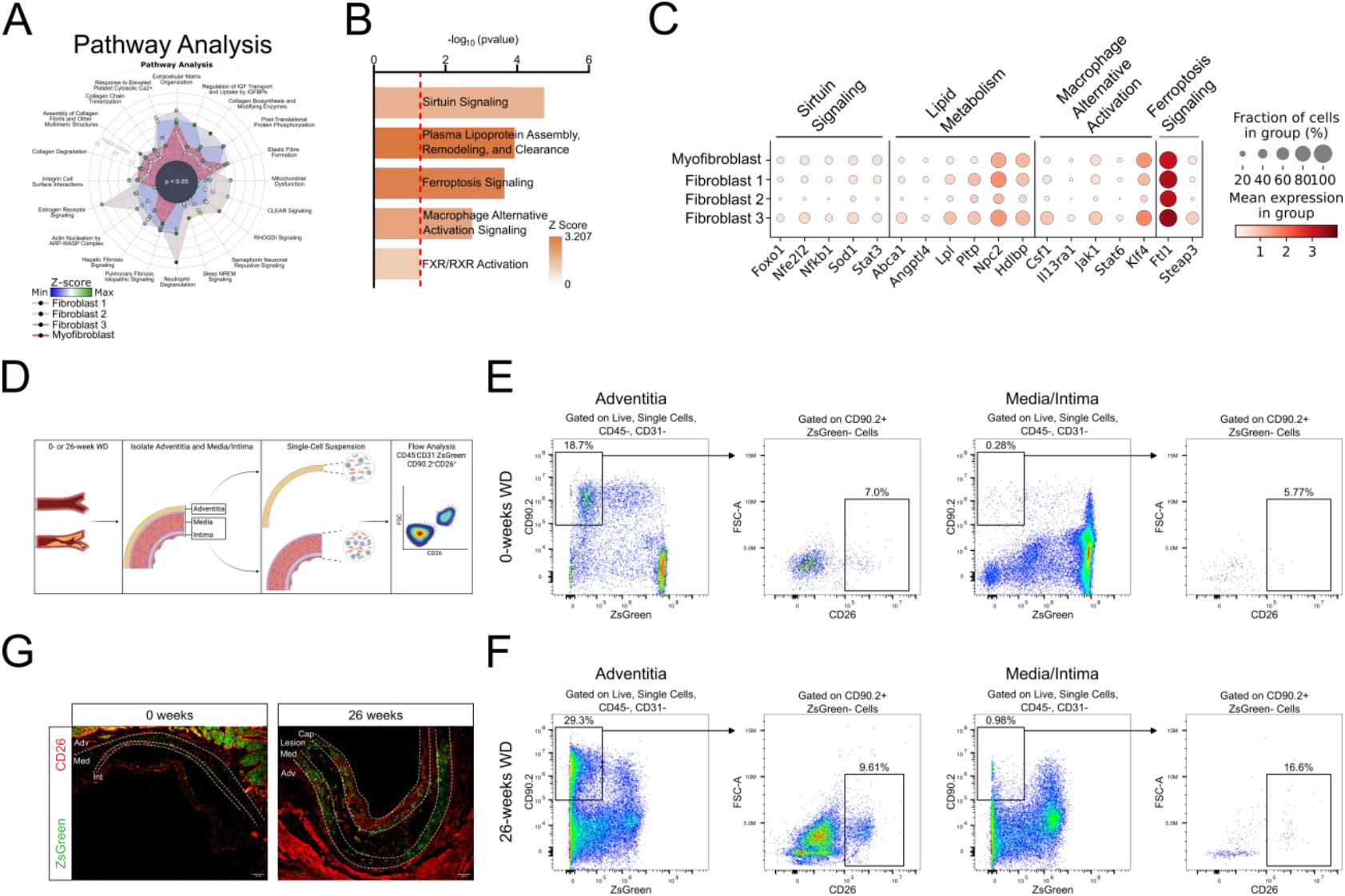
Characterization and Validation of CD26^+^ fibroblast subpopulation. (A) Ingenuity pathway analysis (IPA) using the top DE genes for each fibroblast subpopulation. (B) Selected upregulated pathways in Fibroblast 3 from IPA analysis. (C) Dotplot depicting selected genes that infer functional differences from pathways in panel H. (D and E) Flow cytometry gating strategy used to identify CD26^+^ fibroblasts, which correspond to Fibroblast 3, at 0-(D) and 26-(E) weeks of Western diet feeding. Leukocytes and endothelial cells were first removed by gating on CD45^-^ CD31^-^ cells. Subsequently, CD90.2^+^ZsGreen^-^ cells were selected to isolated purified fibroblasts. Finally, cells were gated on CD26, identifying a clear population of CD26^+^ fibroblasts. (F) Representative immunohistochemistry staining within the adventitia (Adv), media (Med), intima (Int), lesion, and fibrous cap (Cap) of the aortic root of 0- and 26-week Western diet-fed mice. ZsGreen (green) and CD26 (red).

To validate our CITE-seq findings and to determine whether the Fibroblast 3 population resides within both the lesion intima and adventitia during disease progression, we conducted a flow cytometry analysis to measure the proportion of CD26^+^ fibroblasts in the adventitia relative to the media/intima under both healthy conditions and following a 26-week atherogenic diet (**Fig 2D**). We developed a flow cytometry panel using CD26 to discriminate between the fibroblast subsets (**Fig 2E and F**). We excluded CD45^+^ leukocytes and CD31^+^ endothelial cells, ensuring that the remaining cells included only VSMCs and fibroblasts. We then gated on CD90.2^+^ZsGreen^-^ cells, which represent purified fibroblasts. Using flow cytometry expression of CD26, we observed a distinct population corresponding to Fibroblast 3, as identified in our CITE-seq analysis. At 0-weeks of WD feeding the vast majority of fibroblast were in the adventitia, with only approximately 0.3% of cells in the media/intima being fibroblasts, and a smaller fraction being CD26^+^ (**Fig 2E**). However, in advanced atherosclerosis following 26-weeks of WD feeding, almost 1% of cells in the media/intima were fibroblasts, and approximately 17% of these were CD26% (**Fig 2F**). Finally, to complement our flow cytometry analysis and to explore the spatial location of the Fibroblast 3 subpopulation, we performed immunohistochemistry staining to localize CD26^+^ fibroblasts within the arteries of 0- and 26-week WD-fed mice (**Fig 2G**). This analysis revealed the presence of CD26^+^ fibroblasts at both time points, agreeing with our CITE-seq analysis. At 0 weeks, CD26^+^ cells were primarily restricted to the adventitia. However, by 26 weeks a significant proportion of advanced lesions stained positively for CD26 in the intima and fibrous cap, as well as the adventitia, with little to no colocalization with VSMC-derived cells. This suggests that during the progression of atherosclerosis, this fibroblast subpopulation may migrate from the adventitia, where these fibroblasts originate, into the advancing lesion.

## Discussion

In this study, we uncovered substantial phenotypic diversity among fibroblasts within atherosclerotic lesions, identifying three major fibroblast subpopulations and a myofibroblast population that clusters with VSMC-derived chondromyocytes. These findings highlight the overlapping phenotypic characteristics shared by fibroblasts and VSMCs. This represents the first comprehensive multimodal assessment of fibroblasts in atherosclerosis, utilizing advanced single-cell techniques to integrate gene and protein expression data and cell lineage-tracing. Each fibroblast subpopulation exhibited a distinct gene and protein expression profile, highlighting their potential for unique functional roles in the disease process. Among these, we identified for the first time a CD26^+^ fibroblast subpopulation in lesions that increases in lesion intima during disease progression, suggesting that it migrates from adventitia, where it is located in non-disease vessels. The migratory capacity of adventitial fibroblasts toward the lumen has been previously documented in the context of balloon injury^20^, supporting this concept. However, our findings suggest that specific fibroblast subpopulations may exhibit preferential migration during atherosclerosis, adding nuance to this phenomenon. This highlights the potential active role of these cells in shaping the atherosclerotic microenvironment and in modulating features of plaque stability including fibrous cap thickness.

Pathway analysis revealed that this lesional CD26^+^ fibroblast is enriched in genes associated with lipid metabolism, suggesting that it may contribute directly to lipid homeostasis or dysregulation within atherosclerotic plaques. CD26? fibroblasts exhibited significant enrichment in genes linked to macrophage alternative activation, notably Csf1, underscoring their potential role in modulating inflammation and sustaining atherosclerosis through the recruitment, differentiation, and survival of macrophages^21–23^. Additionally, these fibroblasts demonstrated enrichment in sirtuin signaling pathways, which are thought to confer protective effects against atherosclerosis^24^, as well as ferroptosis signaling pathways, which may exacerbate the inflammatory tissue microenvironment and further drive the progression of atherosclerosis^25^. These findings offer significant insights into the heterogeneity of fibroblasts in atherosclerosis, highlighting potential cellular targets for advancing mechanistic studies and developing therapeutic interventions. The interplay between protective and deleterious atherogenic pathways enriched in these fibroblasts likely plays a pivotal role in shaping their contribution to atherosclerotic progression, emphasizing the need to unravel the factors that regulate this balance.

Our research builds upon recent advancements in utilizing single-cell RNA sequencing (scRNA-seq) to explore fibroblast heterogeneity in atherosclerosis, which has predominantly focused on the adventitia^14^. Previous studies identified three distinct fibroblast subtypes with unique developmental trajectories: CD55^+^, CXCL14^+^, and LOX^+^ subtypes. Our work expands on this foundation by identifying a novel CD26^+^ fibroblast subpopulation, which aligns with the previously described CD55^+^ fibroblasts. Notably, while CD26 RNA expression is lower compared to CD55 RNA, CD26 protein is both more specific and significantly more abundant than CD55 protein or any other markers associated with this subpopulation (**Figure 1E and F**), making it a robust marker of this fibroblast subpopulation and one that can be used to isolate these cell by flow sorting for functional and translational studies. Interestingly, while prior studies highlighted Cxcl14^+^ and Lox^+^ as markers for specific fibroblast subpopulations, we were unable to validate their consistent expression or their association with distinct fibroblast subsets in atherosclerotic lesions. This raises questions about the specificity and reproducibility of these markers in the context of atherosclerosis, suggesting potential variability across experimental conditions, models, or stages of disease. Our findings refine the understanding of fibroblast heterogeneity and highlight the need for more robust and reproducible markers to accurately characterize fibroblast subtypes in the complex microenvironment of atherosclerotic plaques.

CD26^+^ fibroblasts have emerged as critical players in a variety of pathological contexts, underscoring their functional importance in both homeostasis and disease. Notably, CD26^+^ scar-forming fibroblasts have been identified in dermal fibrosis, where they are hypothesized to play a pivotal role in modulating the wound-healing process^26^. This association highlights their involvement not only in resolving tissue injury but also in perpetuating fibrosis under pathological conditions. Another study demonstrated that CD26^+^ tissue-resident fibroblasts in breast cancer can transition into inflammatory tumor-associated fibroblasts, promoting tumor cell invasion and recruiting myeloid cells^27^. Adding to their significance, a comprehensive fibroblast atlas across 16 mouse tissues revealed the presence of a CD26^+^ adventitial pan-tissue fibroblast population^15^. This groundbreaking study suggests CD26^+^ fibroblasts constitute a conserved lineage with shared phenotypic and functional characteristics across diverse organ systems. Such findings implicate CD26^+^ fibroblasts in fundamental biological processes, particularly those related to tissue repair, fibrosis, and potentially chronic inflammation. These studies highlight the conserved role of CD26^+^ fibroblasts in tissue maintenance and fibrotic disease pathogenesis. Their consistent presence across tissues and disease models underscores their relevance in conditions involving dysregulated repair mechanisms, such as systemic sclerosis, pulmonary fibrosis, and cardiovascular remodeling. Further characterization of the signaling pathways and interactions mediated by CD26^+^ fibroblasts may inform the development of targeted therapies for fibrotic disorders.

Whether CD26^+^ fibroblasts play roles in fibrous cap formation and plaque stability in atherosclerosis is an important unanswered question. Our immunohistochemistry analysis indicates that this fibroblast subpopulation preferentially localizes to the fibrous cap of lesions, suggesting a potential role in promoting lesion stability. However, this remains speculative, and further investigation using CD26 lineage-tracing models and genetic deletion approaches is necessary to definitely elucidate the origin and role of this fibroblast population in atherosclerotic disease development, progression and stability.

In conclusion, our multi-omic, high-dimensional cell surface phenotyping of fibroblasts in atherosclerosis combined with cell lineage tracing revealed significant lesion fibroblast heterogeneity, including a CD26^+^ subpopulation that likely migrates form adventitia into lesions to contributes to lesion pathophysiology and features of plaque stability. This discovery underscores the potential functional importance of fibroblast subpopulation diversity and functions in the development, progression and complications of atherosclerosis.

## Supporting information

DE Genes

## Acknowledgments

CITE-seq (10x Genomics) was performed in the JP Sulzberger Columbia Genome Center, supported in part through the National Institutes of Health/National Cancer Institute Cancer Center Support Grant P30CA013696. Flow cytometry experiments described in this article were performed in the Columbia Stem Cell Initiative Flow Cytometry core facility at Columbia University Irving Medical Center.

